# Tunable Expression of an AAV Payload Using ADAR-mediated RNA Editing

**DOI:** 10.64898/2026.08.04.742838

**Authors:** Joseph Silverberg, Laura Pereira, Robert Schmidt, Cameron Baptista, Ahil N Ganesh, Nicole Harbaugh, Lauren Moffa, Alexander Metz, Victor Howard, Sean Armour, Daniel M Cohen, Federico Mingozzi

## Abstract

A challenge of “once-and-done” adeno associated virus (AAV)-based gene therapy is the inability to modulate the level of therapeutic protein expression post-administration. Herein, we demonstrate the utility of an adenosine deaminase acting on RNA (ADAR) - mediated gene switch to control AAV-delivered gene expression. Using a premature termination codon (PTC) in the human Factor IX (hFIX) transgene, we established an ON switch, where expression of hFIX is contingent on rescuing the PTC mutation via RNA editing. *In vitro* and *in vivo* studies demonstrated silencing of the hFIX transgene by the PTC mutation and induction of protein expression by administration of an ADAR-recruiting trigger RNA. Mice transduced with a hepatotropic AAV capsid encoding an ApoE-hAAT hFIX-PTC transgene expression cassette showed a dose-dependent response between the levels of LNP-delivered trigger RNA and the amount of plasma hFIX expression achieved. We observed predictable and reproducible levels of hFIX expression upon multiple rounds of RNA editing and demonstrated that this system can achieve clinically relevant levels of hFIX. This work suggests that ADAR-mediated RNA editing may be a valuable tool for tunable expression of therapeutic transgenes in applied gene therapies.

## Introduction

Adeno-associated virus (AAV)-based gene therapies, typically for gene augmentation purposes, result in constitutive and often irreversible gene expression^1,2^. Both the levels of transgene expression and cell type specificity of gene expression are influenced by the choice of vector dose, promoter and capsid serotype^2–5^. Additionally, the introduction of regulatory sequences, such as UTRs, introns, or miRNA binding sites, can further control transgene expression^6,7^. While these approaches have been effective in many clinical settings, strategies that allow for tunable transgene expression after AAV administration would be beneficial in facilitating safer and broader applicability of gene therapies, particularly in the cases in which constant gene expression is unnecessary or potentially harmful.

Numerous options exist that allow for modulation of gene expression levels. For example, inducible promoters offer temporal control, dose-dependent expression, and the potential for increased safety. The tetracycline (Tet) system is one of the most widely studied, with demonstrated efficacy *in vivo* in rodents and non-human primates^8^. However, clinical translation has been hindered by concerns over immunogenicity due to its bacterial origin^8,9^. Similarly, the rapamycin-inducible system offers transcriptional control using an FDA-approved drug, but its use is complicated by potential interactions with the endogenous mTORC1 pathway, which can affect cellular metabolism and proliferation, as well as by slow off-kinetics following drug withdrawal^10,11^. Alternatively, regulating protein levels using degrons has several benefits such as the precise control of protein stability and the ability to quickly degrade unwanted proteins which can be vital in therapeutic applications^12^. Despite these advantages, there are challenges with this application including off-target effects and difficulty in obtaining efficient and specific exposure of the degrons to the cellular degradation machinery. Recently, approaches that involve RNAi-based regulation and microRNA antagonists (REVERSIR™) or the employment of drug-responsive RNA aptamers are emerging as valuable tools to tune the expression of a therapeutic transgene post-dosing and efforts are underway to test their safety and efficacy in AAV cassettes^13–15^.

Gene editing technologies further expand the toolkit to control AAV-delivered transgene expression post-dosing. Notably, the endogenous adenosine deaminase acting on RNA (ADAR) enzymes, which are ubiquitously expressed across mammalian tissues, have shown promise for therapeutic RNA editing^16,17^. ADARs can be recruited to correct endogenous mutations by editing adenosines to inosines (read as guanosine by the tRNA) within target double stranded RNAs (dsRNA)^18^. Here, we present a novel strategy for post-delivery control of AAV transgene expression using ADAR-mediated RNA editing. By delivering a trigger RNA via LNPs, we successfully recruited endogenous ADAR to convert a premature termination codon (PTC) to a functional amino acid codon in the AAV-delivered mRNA transgene, effectively acting as an ON switch. Transgene expression went from undetectable to peaking within 24 hours of trigger RNA dosing. In our system, re-dosing the trigger RNA did not diminish the efficacy of transgene induction. This new approach provides a tunable, non-immunogenic, and reversible method for controlling transgene expression *in vivo*, offering a new avenue for safer and more flexible gene therapies.

### Results

To control gene expression through ADAR-based RNA editing, we introduced a premature TAG amber stop codon in lieu of a tryptophan codon (TGG) within Exon 2 of hFIX in a pAAV plasmid to generate hFIX-PTC (Figure 1A). This mRNA template is amenable to ADAR-mediated reversion of the PTC by conversion of adenosine to inosine (inosine is decoded by tRNAs as guanosine)^19^. In transfection experiments, plasmids carrying the PTC mutation at residue W118 showed undetectable levels of the protein as assessed by automated Western blot assay (middle lane, Figure 1B). Replacement of other tryptophan codons by PTC mutations yielded variable levels of hFIX expression in the same assay (Figure S1). This data showed the importance of the empirical selection of the PTC location within the mRNA and justified using the W118, referred to as hFIX-PTC for subsequent experiments.

**Figure 1.**
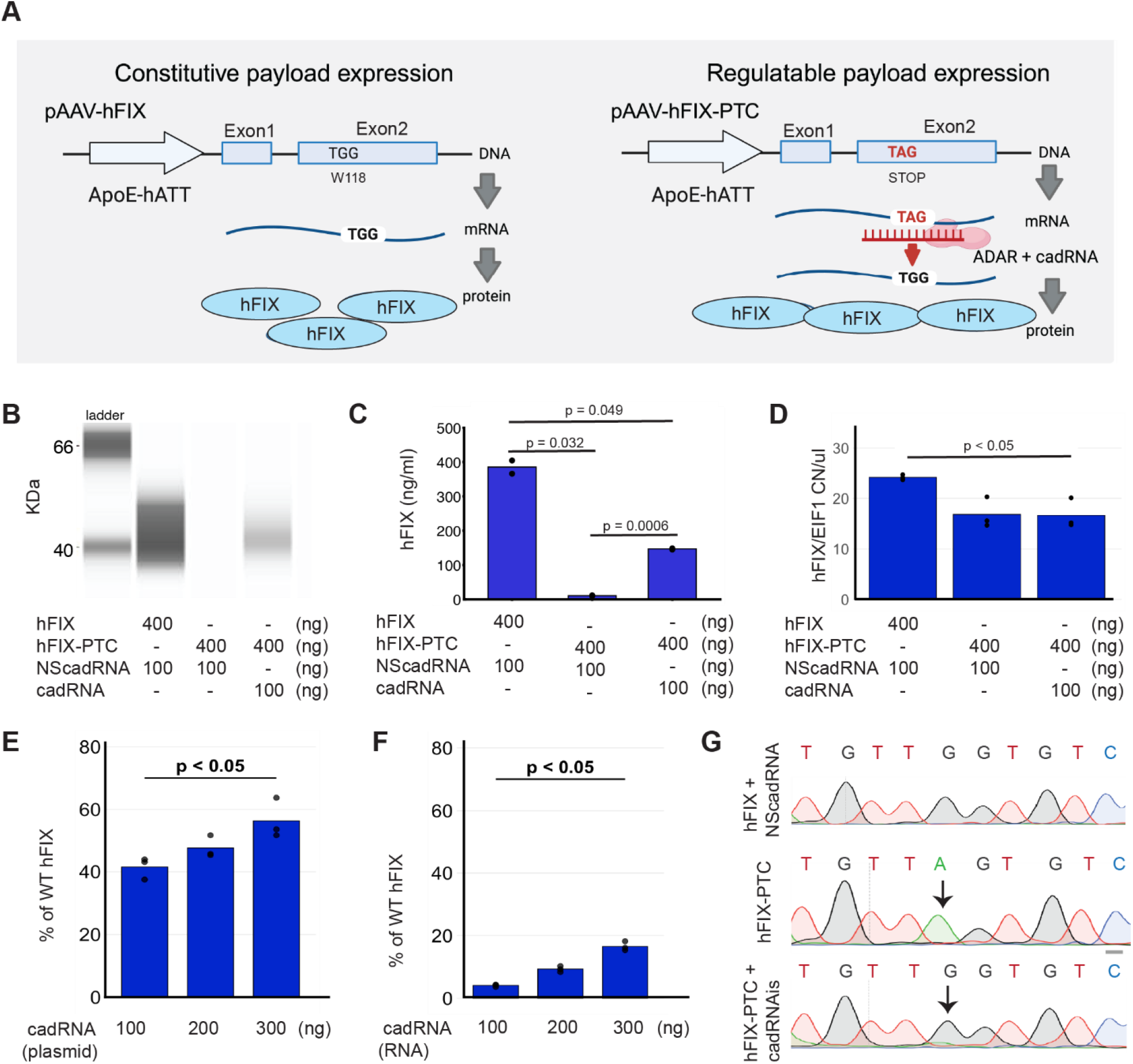
In vitro proof-of-concept for endogenous ADAR-mediated rescue of hFIX-PTC. **(A)** Schematic representation of the constitutive wild-type hFIX compared to the hFIX-PTC regulatable system. Functional protein expression is contingent upon ADAR-mediated A-to-I editing to revert the premature termination codon (TAG) to a tryptophan codon (TGG). **(B)** Western Blot showing no signal from hFIX-PTC and partial rescue of hFIX protein in Huh7 cells following co-transfection with site-specific trigger RNA in circular form (cadRNA). NS: non-specific control cadRNA. **(C)** Quantitative analysis by ELISA of secreted hFIX protein in the same samples. Data demonstrate significant protein recovery in cadRNA-treated cells compared to PTC controls, p = 0.0006 by Welch’s t-test (n=2). **(D)** hFIX mRNA levels measured by RT-qPCR normalized to the EIF1 housekeeping gene and expressed as copy number per ul of cDNA. While the PTC mutation significantly reduced mRNA abundance, cadRNA treatment did not significantly alter transcript levels, p < 0.01 by One-way ANOVA with Tukey’s post-hoc test (n=3). **(E)** Dose-dependent increase in hFIX protein expression following transfection of increasing concentrations of plasmid-encoded cadRNA in Huh7 cells, expressed as a percentage of wild-type hFIX levels. **(F)** hFIX protein expression with increasing amounts of *in vitro* transcribed (IVT) circular RNA, expressed as a percentage of wild-type hFIX levels. (**G)** Molecular validation of RNA Editing by Sanger sequencing chromatograms of cDNA derived from hFIX, hFIX-PTC and hFIX-PTC co-transfected with cadRNA in plasmid form. The arrow denotes the targeted A-to-I (read as G) editing event at codon 118.

To test the ability of endogenous ADAR enzymes to edit a premature stop codon in a model gene of interest, we transiently transfected cells with plasmids encoding an ApoE-hAAT driven hFIX-PTC gene and a U6 promoter-driven circular ADAR-recruiting RNA (cadRNA) featuring interspersed loops^20^. Forty-eight hours post-transfection, robust hFIX protein expression was detected via Western blotting only in cells receiving the site-specific cadRNA (targeting nucleotide 353), whereas expression was absent in the non-specific control (NScadRNA) (Figure 1B). ELISA assays confirmed that while the PTC mutation suppressed hFIX expression relative to wild-type, the addition of cadRNA mediated a significant recovery, restoring hFIX protein to approximately 38% of wild-type levels (Figure 1C).

Interestingly, RT-qPCR analysis revealed a significant reduction of hFIX mRNA of around 30% in the presence of the PTC, likely due to nonsense-mediated decay (NMD). Addition of the cadRNA did not alter total mRNA levels relative to the NScadRNA control (Figure 1D). This observation is consistent with ADAR-mediated induction of FIX expression through translational read-through of the PTC rather than mRNA stabilization. Dose-escalation studies confirmed a linear, dose-dependent induction of hFIX protein (Figure 1E-F), with plasmid-based delivery of a cadRNA achieving superior results compared to synthetic circular RNA (60% vs. 16% of wild-type levels, respectively). Site-specific adenosine-to-inosine (A-to-I) editing at codon 118 was confirmed via Sanger sequencing. Although mRNA editing was efficient, the persistence of a minor unedited adenine peak (Figure 1G) suggests that incomplete protein rescue may be driven by a subset of unedited FIX transcripts. While Sanger chromatograms qualitatively confirm the presence of edited species, they are not strictly quantitative for precise nucleotide ratios. Notably, no collateral ‘off-target’ editing was observed within the target amplicon (data not shown).

Next, we focused on evaluating the potential of the ADAR-based editing to regulate AAV-mediated gene expression *in vivo*. Mice were administered a hepatotropic AAV (AAV Spark-100) encoding either hFIX (n=4; positive control) or hFIX-PTC (n=24; ADAR-mediated switch) (Figure 2A). While the control group received a therapeutically relevant dose of 5E9 vector genomes (vg), the hFIX-PTC cohort was dosed an order of magnitude higher (5E10 vg) to compensate for previously observed NMD-mediated transcript degradation. This higher dose ensured sufficient mRNA substrate for subsequent ADAR-mediated editing. Two weeks post-AAV administration, plasma hFIX levels in the control cohort reached 80 ng/mL, whereas hFIX remained undetectable in the hFIX-PTC group (Figure 2B). These *in vivo* results corroborate our previous *in vitro* findings, confirming that the presence of the PTC effectively abrogates hFIX protein expression.

**Figure 2.**
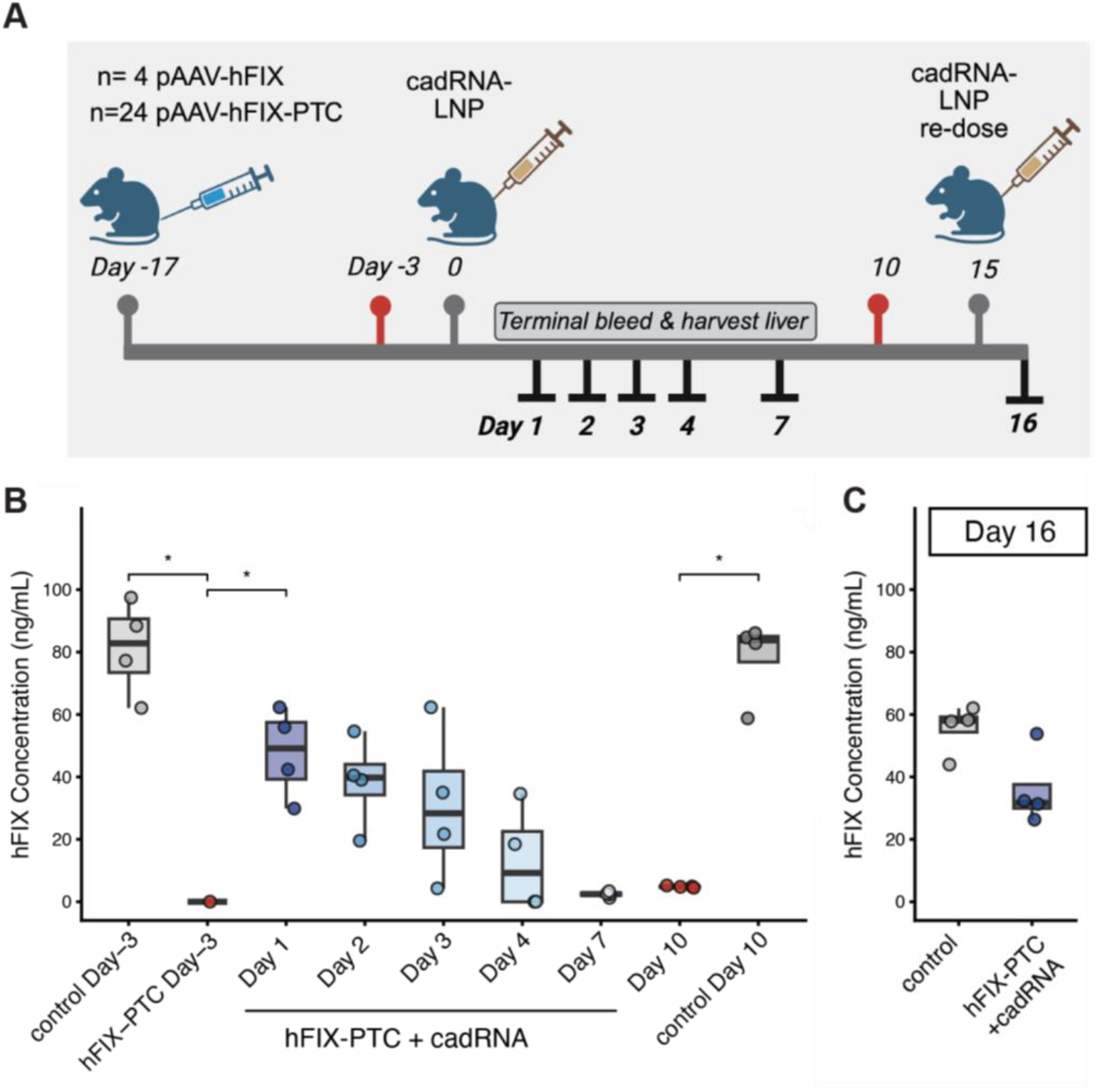
Determining the therapeutic window of cadRNA induced hFIX expression via endogenous ADAR. **(A)** Timeline of mouse proof-of-technical principle study using an ADAR-tunable AAV. A control group was dosed with pAAV-hFIX while 6 additional cohorts were dosed with pAAV-hFIX-PTC (Day −17). Two weeks post-AAV dosing, blood samples were taken for ELISA analysis of hFIX levels (Day −3 samples). Following cadRNA-LNP administration on Day 0, pAAV-hFIX-PTC cohorts were sacrificed on different days, and plasma and liver tissue collected. On Day 10, blood samples were collected from the control and remaining pAAV-hFIX-PTC cohort. This single cohort was redosed with cadRNA-LNP on Day 15 and sacrificed on Day 16. **(B)** Plasma hFIX expression levels measured by ELISA as a function of time post cadRNA-LNP delivery. Boxplots in gray show the range of hFIX observed in the hFIX control animals (Day −3 and Day 10). Plasma hFIX levels were not statistically significant across successive days in the time course (e.g. D1 vs D2 post cadRNA-LNP dosing - run statistics since this was Joe’s statement). **(C)** Mice redosed with cadRNA-LNP at Day 15 were sacrificed at Day 16, and the induction of plasma hFIX were measured by ELISA in these animals as well as in the control group.

Two weeks post-AAV administration, mice were dosed with lipid nanoparticles (LNPs) encapsulating trigger RNA payloads. To measure delivery efficiency in the mouse liver, the trigger cadRNA loaded LNPs were co-formulated with a *Gaussia* luciferase reporter mRNA at a 4:1 mass ratio and dosed at 5 µg of mRNA per animal. In order to capture the kinetic profile of the RNA editing response, study cohorts were sacrificed at days 1, 2, 3, 4, or 7 post-LNP administration (Figure 2A). Successful hepatic delivery was confirmed by robust plasma luciferase activity on day 1, followed by a characteristic exponential decay (Figure S2A). Within 24 hours of trigger-LNP delivery, we observed a profound spike in plasma hFIX, peaking at an average of 48 ng/mL, or around 60% of the control levels. Following this single dose, hFIX levels declined to 38.5 ng/mL at day 2, 30.8 ng/mL at day 3, 13.3 ng/mL at day 4, 5ng/mL at day 7 and fell below the limit of quantitation by day 10 (not a terminal bleed). To assess the feasibility of redosing, the animal cohort that showed no hFIX expression at day 10 received a second LNP dose on day 15, which successfully yielded a reproducible hFIX spike of 35 ng/mL, or around 60% of the control levels measured at the same time point. This indicates that repeat RNA editing is feasible and produces predictable transgene expression profiles. Finally, to evaluate whether the ADAR-responsive PTC mutation affected either transduction or stability of the resultant mRNA transcript, we extracted vector genomes and mRNA from the liver of the transduced mice at the end of the study. Transcripts and vector genomes were measured by qPCR using absolute quantification. The hFIX-PTC transcript abundance per AAV vector genome copy number (VGCN) were statistically indistinguishable from wild-type hFIX mRNA transcripts abundance even though hFIX-PTC was dosed with one order of magnitude more virions (Figure S2B). The lack of a statistically significant difference in mRNA:VGCN ratio for the wild-type FIX vectors versus the PTC mutant suggests that, in contrast to the in vitro system, the stability of the FIX transcript does not appear to be sensitive to the presence of the PTC mutation.

Given the transient duration and incomplete rescue of FIX expression by ADAR *in vivo*, we explored whether ADAR efficiency and hFIX expression durability could be boosted by higher doses of either the trigger RNA or by boosting ADAR levels. When mice were dosed with increasing amounts of cadRNA, we noted a dose-dependent response between the amount of cadRNA LNPs delivered and the level of hFIX achieved, measured at different timepoints after LNP dosing (Figure 3A). At the highest dose, 15 µg of trigger cadRNA, hFIX levels rose to 400 ng/mL, whereas at a dose of 3.75 µg of trigger cadRNA modest levels of plasma hFIX were achieved (31 ng/mL) (Figure 3A). These results showed that the trigger cadRNA and not the hFIX-PTC mRNA substrate is the limiting factor in the ADAR-editing reaction. Similarly to our previous data, hFIX expression levels decreased by around 20% percent from Day 1 to Day 3. At Day 9 there was no detectable hFIX in the mice dosed with the cadRNA-LNP regardless of the dose. We therefore infer that the durability of the response may be determined by the half-life of the trigger cadRNA.

**Figure 3.**
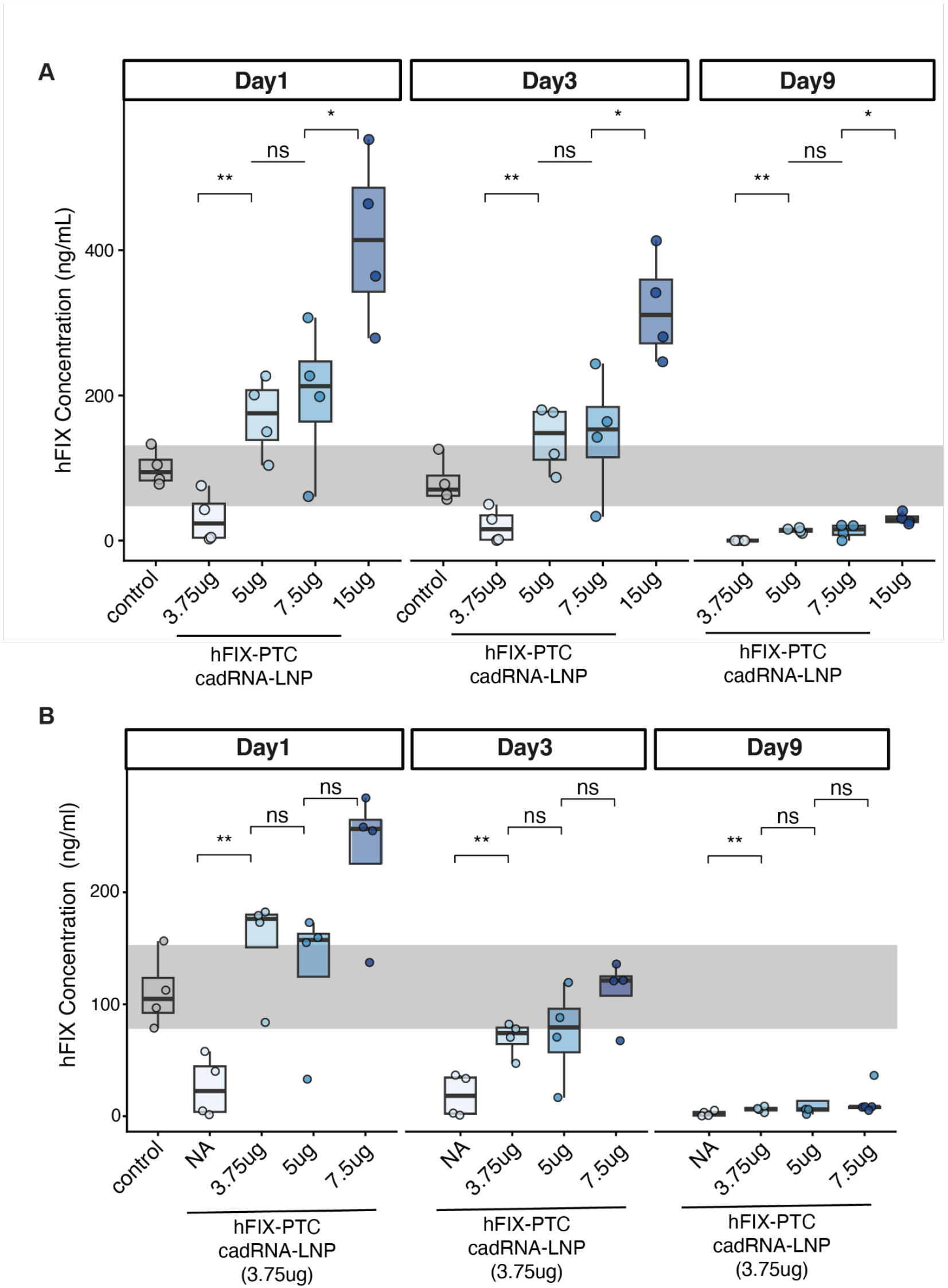
ADAR-mediated rescue of the PTC is dependent on trigger RNA dose and LNP formulation *in vivo*. **A**. Dose escalation of the trigger cadRNA-LNP boosted hFIX to supraphysiological levels. The grey bar represents the range of hFIX expression in the animals dosed with pAAV-hFIX (control cohort). **B**. Dose escalation of ADAR mRNA and its effect on ADAR editing. Dose 1; no exogenous hADAR mRNA was added. *, p < 0.05.

Next, we tested if supplying exogenous hADAR (delivered as mRNA) could increase the efficacy or durability of the ADAR-based mRNA editing. When delivering a constant cadRNA dose (3.75ug) and increasing amounts of hADAR mRNA, we observed a transient increase in hFIX (Figure 3B) that was statistically significant at Day 1. Higher dose of ADAR mRNA did not increase the response, indicating that this effect saturates at the 3.75ug mRNA dose. Together, these data highlight the importance of trigger RNA dose and half-life of the trigger RNA as rate-limiting factors for efficient ADAR-mediated editing in mouse liver.

## Discussion

The ability to modulate transgene expression from an episomal vector post-administration represents a critical evolution in the safety paradigm of AAV-mediated gene therapy. While ‘once-and-done’ approaches have revolutionized the treatment of monogenic disorders, their constitutive nature offers no recourse for patients experiencing adverse immune responses or protein levels outside the optimal therapeutic window where chronic over-expression can lead to undesired phenotypes. Our novel ADAR-mediated switch addresses this by providing a turn-on solution; the therapeutic gene remains dormant due to the engineered amber STOP codon until activated by a transient trigger cadRNA that edits it to a tryptophan. Unlike traditional Tet-ON or rapamycin-inducible systems that rely on potentially immunogenic transactivators, our approach utilizes endogenous ADAR enzymes to minimize metabolic and off-target interference. While other ADAR-based strategies utilize AAV-delivered guide RNAs for permanent CNS-targeted editing^20^, our LNP-delivered trigger provides the unique advantage of temporal, reversible control offering a more flexible and clinically viable path for precise gene regulation.

Our *in vivo* results from mice studies demonstrate a versatile dynamic range for hFIX expression, suggesting that this platform can be tailored to meet specific patient physiological needs through varied dosing strategies. We observed that hFIX protein levels could be titrated from a baseline below the limit of quantitation to therapeutically relevant concentrations of 400 ng/mL, depending on the cadRNA-LNP dose. This wide therapeutic window enables the delivery of AAV vectors expressing proteins that would need to be tightly regulated. It would also enable novel clinical paradigms like a hybrid dosing approach. In such a scenario, a patient could be administered a conservative dose of AAV expressing the wild-type sequence of a protein to establish a low-level, constitutive baseline expression. A second AAV vector expressing a regulatable version of the protein could be administered to drive on-demand spikes of expression. This regulatable component would remain inactive until a “boost” is required at which point an LNP-trigger could be administered to acutely elevate protein levels. The ability to achieve predictable and reproducible induction upon re-dosing, as seen in our Day 15 challenge, confirms that this system can function as a “biologic rheostat,” allowing to provide “on-demand” therapeutic spikes within a safe and controlled homeostatic background.

Despite the versatility of this platform, several technical considerations remain to be addressed. First, the current study utilized liver-targeted LNPs, restricting the scope of this inducible switch to hepatic applications. However, as the field of extra-hepatic delivery evolves, the ubiquitous expression of endogenous ADAR enzymes suggests that this system could be relevant to other target organs, such as the central nervous system or retina, by utilizing tissue-specific LNP formulations. Second, we observed that the duration of transgene induction is transient, likely governed by the *in vivo* half-life of the trigger cadRNA. Further optimization of cadRNA stability perhaps through advanced chemical modifications or structural tuning may be required for applications necessitating more prolonged expression. Finally, our *in vitro* screening revealed that the placement of the PTC is a critical determinant of system performance; not all Trp-to-Stop substitutions effectively blocked FIX protein production. The need to screen multiple PTC candidates was not wholly unexpected, given the known phenomenon that PTCs are more permissive to ribosomal read-through than native termination codons, and that neighboring codon context has been shown to impact the termination efficiency of PTCs^21^. Better understanding of the factors governing the termination efficiency of PTCs is likely to inform the application of ADAR-responsive PTCs for use in regulatable gene therapies.

In conclusion, we created an expression switch that can regulate an AAV-delivered transgene. This expression switch activates the production of protein from a dormant state due to RNA editing occurring from the cadRNA ability to recruit endogenous ADAR to edit the PTC to the native W codon. Our results demonstrated RNA editing *in vitro* and *in vivo* of the hFIX transgene delivered via an AAV, and this could enable tunable control of a therapeutic transgene at physiologically-relevant levels.

## Materials and Methods

### Premature termination codon (PTC) Construct Design

Tryptophan (W) amino acids that could be changed to a stop codon such as TAG or TGA and undergo editing by ADAR were identified within the coding sequence of the hFIX transgene utilizing the Geneious Prime software (Dotmatics, Auckland, New Zealand). We identified four codons at amino acid (AA) 88AA, 118AA, 240AA, and 261AA, all of which could be changed to a stop codon. W at positions 118AA and 240AA were changed to TAG stop codon, while W at 88AA and 261AA were changed to TGA stop codon. All W AA changes can be targeted by ADAR editing. Q5® Site-Directed Mutagenesis Kit (NEB, Cat: E0554S) was used to generate the stop codons in the hFIX transgene. Primers to conduct the site-directed mutagenesis for each AA were designed using the NEBaseChanger Tool (Ipswich, MA), and oligos were synthesized by Integrated DNA Technologies (Coralville, Iowa). The manufacturer’s protocol for the Q5® Site-Directed Mutagenesis Kit was followed to generate the pAAV-hFIX with the different stop codons. The PCR product was then transformed into One Shot™ Stbl3™ Chemically Competent E. coli (Invitrogen, Cat: C737303) following the protocol supplied. Colonies were picked and grown overnight in 5mL cultures of LB broth with kanamycin in a shaking incubator at 30°C. DNA was isolated the following day using the QIAprep Spin Miniprep Kit (Qiagen, Cat: 27106) as outlined in the respective protocol. Plasmids were confirmed correct by NGS sequencing performed by Plasmidsaurus (South San Francisco, CA).

### cadRNA and LNP Formulations

The interspersed cadRNA was designed utilizing the approach laid out by Katrekar et al. with the 200bp trigger RNA length, the edit at the 100bp mark, and interspersed loops on both ends of the sequence^20^. The cadRNA is driven from the human U6 promoter and circularized by including the twister ribozyme regions on the 5’ and 3’ ends. Sequences were designed with Geneious Prime software (Dotmatics, Auckland, New Zealand) and synthesized into pUC57 at Azenta (South Plainfield, NJ). Circular mRNA for the cadRNA was synthesized at GenScript (Piscataway, NJ), and control Gaussia Luciferase mRNA was synthesized at TriLink BioTechnologies (San Diego, CA). Both the cadRNA and Gaussia Luciferase mRNA were formulated into separate proprietary LNPs that are liver-targeted and then mixed. LNP formulation was conducted in-house following in-house generated encapsulation protocols utilizing proprietary components.

### Cell Culture and Transfection

In vitro experiments were conducted in Huh7 cells grown in DMEM supplemented with 10% FBS in an incubator at 37°C with a 5% CO_2_ atmosphere. Huh7 cells were seeded into a 24-well plate (Corning, Cat: 3527) at a density of 1.00E+5 cells/well. Cells were transiently transfected with 500ng of total nucleic acids. Transfection reagent Lipofectamine 3000(Invitrogen, Cat: L3000001) was used for all DNA-based transfections following the manufacturer’s protocol. Transfection reagent TransIT-mRNA Transfection Kit (MirusBio, Cat: MIR 2225) was used for all RNA-based transfections following the manufacture protocol. Media and cells were harvested at 48h after transfection.

### Detection of FIX Antigen and Protein

An ELISA to detect the hFIX antigen from cell supernatant or mouse plasma was conducted with Factor IX Paired Antibody Set (Affinity Biological, Cat: FIX-EIA), and an in-house protocol for a sandwich ELISA was followed. The ELISA was read via BioTek Synergy H1 Multimode Reader (Agilent, Santa Clara, CA USA) and analyzed via SoftMax Pro Software (Molecular Devices, LLC. San Jose, CA USA). Data was graphed via GraphPad Prism 10 software (GraphPad, San Jose, CA).

A capillary-based western blot analysis (WES) (Protein Simple, San Jose, CA) was conducted to detect the FIX protein with 12-230 kDA Wes Separation Module (Protein Simple, Cat: SM-W004) following the manufacturer’s protocol. The primary antibody for the detection of hFIX was the capture antibody in the Factor IX Paired Antibody Set (Affinity Biological, Cat: FIX-EIA) used at a 1:100 dilution. The secondary antibody was the detection antibody in the Factor IX Paired Antibody Set (Affinity Biological, Cat: FIX-EIA) used at a 1:100 dilution. Data was analyzed using the software Compass (Protein Simple, San Jose, CA).

### AAV Vector

AAV-hFIX and AAV-hFIX-PTC were prepared in-house by our Capsid Engineering Team at Spark Therapeutics now Roche Innovation Center Philadelphia with an in-house protocol for AAV production which utilized a triple transfection and purification via cesium chloride gradient. Titers for AAV vectors were determined with a RT-qPCR based assay using TaqMan™ Universal PCR Master Mix (Applied Biosystems, Cat: 4304437) and following the manufacturer’s protocol. The vectors were aliquoted and stored at −80°C before use.

### Animal Experiment

Eight week-old male C57BL/6J mice were acquired from Jackson Laboratories and group housed in controlled temperature with ∼55% humidity under a 12h light/dark cycle with enrichment. AAVs were injected intravenously at a dose of 5.00E+09 or 5.00E+10 vector genomes per mouse. LNPs were injected intravenously at doses of 0.3mpk, 0.45mpk, 0.6mpk, and 0.75mpk. Plasma and liver tissue samples were collected at the indicated time points for each study. When indicated, liver tissue samples were also collected at the study endpoint. All in vivo procedures were approved by the Institutional Animal Care and Use Committee (Lampire Biologic Labs, Pipersville, PA).

### Luciferase Assay

Gaussia Luciferase was detected with the Renilla Luciferase Assay System (Promega, Cat: E2820) following the manufacturer’s protocol. Mouse plasma was diluted 1:10 into control C57BL/6 plasma and plated into a 96-well black plate with a solid bottom. The plate was then loaded into GloMax Discover (Promega, Madison, WI), and 100µL of assay buffer was injected one well at a time at a speed of 200µl/s with a luminescence kinetics integration time of 1 second. Relative luminesce units are graphed via GraphPad Prism 10 software (GraphPad, San Jose, CA).

### Cell and Tissue Extraction of RNA and Genomic DNA

RNA from cells and liver tissues were extracted using RNeasy Plus Mini Kit (Qiagen, Cat: 74134) following the manufacturer’s protocol. cDNA was synthesized with a High-Capacity cDNA Reverse Transcription Kit with RNase Inhibitor (Applied Biosystems, Cat: 4374967) following the manufacturer’s protocol. 500ng of cDNA was amplified by PCR by NEBNext® High-Fidelity 2X PCR Master Mix (NEB, Cat: M0541L) with primers specific to the target site (FWD: 5’-TGAGGAAGCCAGAGAAGTGT-3’ / REV:5’-CTCAGTGCAGCTACAGACCACT-3’), generating a PCR product of ∼500bp following the manufacturer’s protocol. Following the manufacturer’s protocol, The PCR product was purified with Monarch® Spin PCR & DNA Cleanup Kit (NEB, Cat: T1135). The purified PCR product was premixed with a sequencing primer (5’-GGATGGTGATCAATGTGAGAGC-3’) upstream of the edit site and sent for Sanger Sequencing at Azenta (South Plainfield, NJ). Trace files were analyzed in SnapGene software for the presence of editing.

Genomic DNA was extracted from liver tissues using the QIAamp DNA Mini Kit (Qiagen, Cat: 51304) following the manufacturer’s protocol.

### Detection of hFIX mRNA and Vector Genome Copy Number (VGCN)

2µL of cDNA or gDNA were used to set up a 20µL RT-qPCR reaction using TaqMan™ Universal PCR Master Mix (Applied Biosystems, Cat: 4304437) and following the manufacturer’s protocol. Primers and probe assays were designed to detect hFIX and housekeeping genes EIF1 in humans and mice. They were ordered from IDT as a PrimeTime™ qPCR Probe Assay with 5’ FAM reporter dye and 3’ZEN/IowaBlackFQ quencher. Three technical replicates were done for each sample. RT-qPCR was run on the QuantStudio 7 (ThermoFisher, Waltham, MA) with standard curves. Samples were analyzed in the QuantStudio software ExpressionSuite (ThermoFisher, Waltham, MA), where mRNA and VGCN were normalized to the housekeeping genes h/mRLPL0 and graphed in GraphPad Prism 10 software (GraphPad, San Jose, CA).

hFIX primer-F(ORL_131): 5’-CCTGCTCTGCTCGAATGTA-3’,

hFIX primer-R(ORL_132): 5’-TACACTCTCTCCAGATTCCC-3’,

hFIX Probe (FIX qPCR P2): 56-FAM/ACC CAA AGA G/ZEN/GTA TAA CTC TGG CAA GCT/3IABkFQ/,FIX qPCR gBlock (IDT, Ref 580576552; 150 bp)

Housekeeping primer-F (EIF1-1 F): 5’-GAAACGGCAGGAAGAGCCCTTA-3’, Housekeeping primer-R (EIF1-1 R): 5’-CGGATGCTCAATTACAGTACCAT-3’,

Housekeeping probe (EIF1-1 Probe): ACTGTC CAAGGGATCGCTGATGA (primerbank ID: 77404355c1), EIF1-1 gBlock (IDT; 119 bp)

BGHpA primer 1: 5’-CTTGCCTTCCTTGACCCT-3’, BGHpA primer 2: 5’-CCCAGAATAGAATGACACCTACT-3’, BGHpA probe: 56-FAM/5’-TTAGGAAAG-3’/ZEN/5’-GACAGTGGGAGTGGC-3’/3IABkFQ

## Appendix

**Figure S1.**
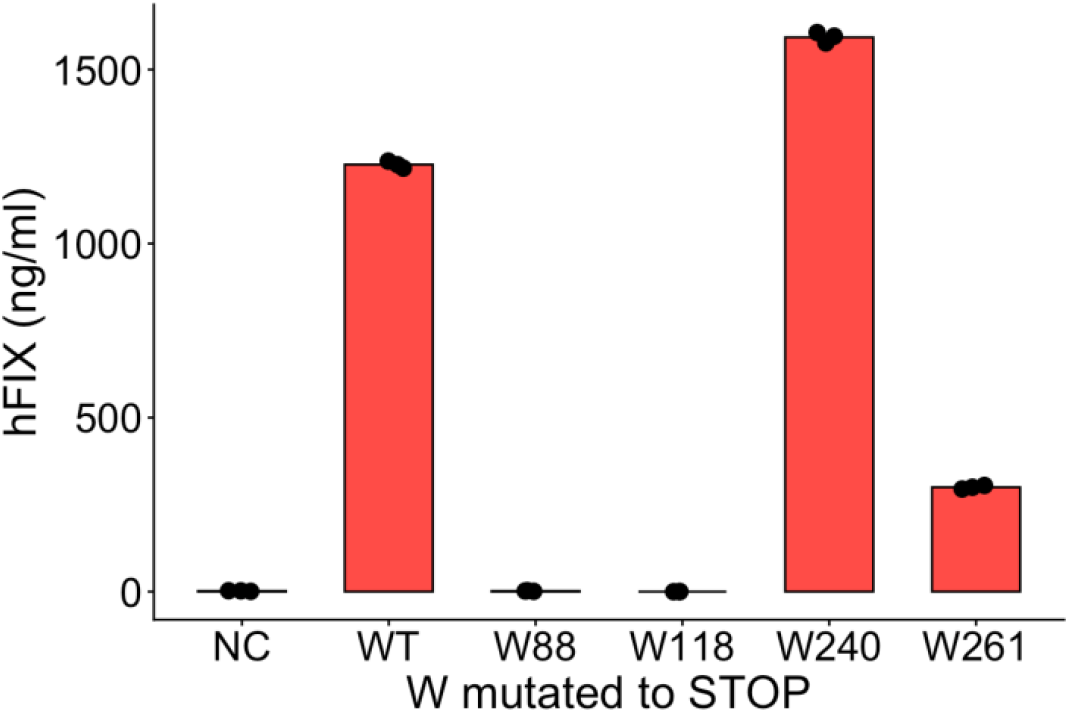
ELISA showing FIX expression levels obtained from Huh7 transiently transfected cells with plasmids of the PTC codons at different AA locations within the FIX transgene. PTC at positions 240AA, and 261AA demonstrated leaky expression.

**Figure S2.**
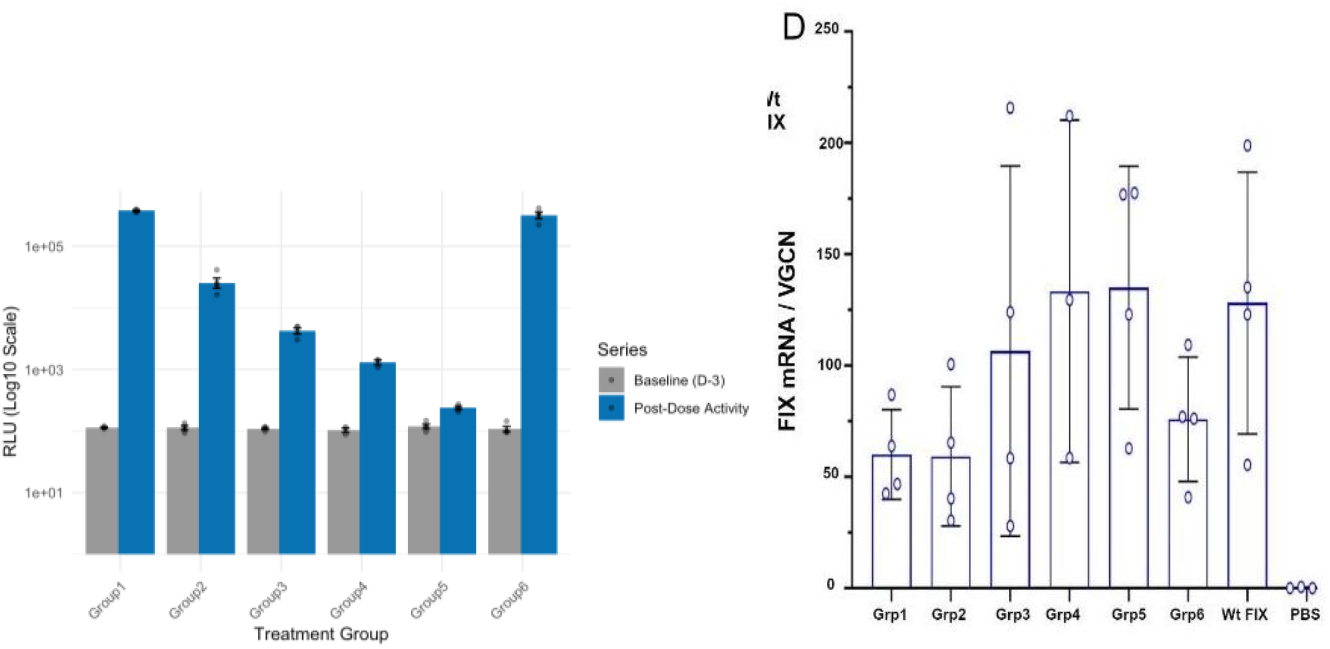
A. Gaussia luciferase was co-delivered with trigger RNA to assess LNP liver tropism. Luciferase activity demonstrated successful LNP-mediated liver delivery of RNA. Gaussia luciferase activity in plasma was undetectable in every group at Day −3 (pre-treatment bleeds). Gaussia luciferase was measured in the terminal bleed for Group 1 (Day 1), Group 2 (Day 2), Group 3 (Day 3), Group 4 (Day 4), Group 5 (Day 7) and Group 6 (Day 16, redose) and had an observed half-life of ∼6hrs with first-order exponential decay kinetics, R^2^ = 0.996. B. Normalized FIX transcript abundance per AAV vector genome copy showed roughly comparable levels of transduction and gene expression at the mRNA level. No statistical differences were observed for normalized FIX mRNA/VGCN ratio across study groups.

